# Compression of short-read sequences using path encoding

**DOI:** 10.1101/006551

**Authors:** Carl Kingsford, Rob Patro

## Abstract

Storing, transmitting, and archiving the amount of data produced by next generation sequencing is becoming a significant computational burden. For example, large-scale RNA-seq meta-analyses may now routinely process tens of terabytes of sequence. We present here an approach to biological sequence compression that reduces the difficulty associated with managing the data produced by large-scale transcriptome sequencing. Our approach offers a new direction by sitting between pure reference-based compression and reference-free compression and combines much of the benefit of reference-based approaches with the flexibility of *de novo* encoding. Our method, called path encoding, draws a connection between storing paths in de Bruijn graphs — a common task in genome assembly — and context-dependent arithmetic coding. Supporting this method is a system, called a bit tree, to compactly store sets of kmers that is of independent interest. Using these techniques, we are able to encode RNA-seq reads using 3% – 11% of the space of the sequence in raw FASTA files, which is on average more than 34% smaller than recent competing approaches. We also show that even if the reference is very poorly matched to the reads that are being encoded, good compression can still be achieved.

## 1. INTRODUCTION

The size of short-read sequence collections is often a stumbling block to rapid analysis. Central repositories such as the NIH Short Read Archive (SRA; National Institutes of Health, 2014) are enormous and rapidly growing. The SRA now contains 2.5 petabases of DNA and RNA sequence information, and due to its size, it cannot be downloaded in its entirety by anyone except those with enormous resources. When select experiments are downloaded, the local storage burden can be high, limiting large-scale analysis to those with large computing resources available. Use of cloud computing also suffers from the data size problem: often transmitting the data to the cloud cluster represents a significant fraction of the cost. Data sizes also hamper collaboration between researchers at different institutions, where shipping hard disks is still a reasonable mode of transmission. Local storage costs inhibit preservation of source data necessary for reproducibility of published results.

Compression techniques that are specialized to short-read sequence data can help to ameliorate some of these difficulties. If data sizes can be made smaller without loss of information, transmission and storage costs will correspondingly decrease. While general compression is a long-studied field, biological sequence compression — though studied somewhat before short-read sequencing (e.g. Ladner, 2004; Matsumoto et al., 2000) — is still a young field that has become more crucial as data sizes have outpaced increases in storage capacities. In order to achieve compression beyond what standard compressors can achieve, a compression approach must be tailored to the specific data type, and it is likely that different compression approaches are warranted even for different short-read experimental settings such as metagenomic, RNA-seq, or genome assembly applications.

Here, we present a new compression algorithm for collections of RNA-seq reads that outperforms existing compression schemes. RNA-seq experiments are extremely common, and because they are repeated for many different conditions, the number of future experiments is nearly unbounded. Among the SRA’s 2,500 terabases, there are 72,304 experiments labeled “RNA-seq” that contain short-read sequences of expressed transcripts. While the compression technique we describe here was motivated by, and optimized for, RNA-seq data, it will work for any type of short read data.

Existing short-read compression approaches generally fall into categories: reference-based schemes (Campagne et al., 2013; Fritz et al., 2011; Li et al., 2014) attempt to compress reads by aligning them to one or more known reference sequences and recording edits between the read and its mapped location in the reference. *De novo* compression schemes (Adjeroh et al., 2002; Bhola et al., 2011; Bonfield and Mahoney, 2013; Brandon et al., 2009; Burriesci et al., 2012; Cox et al., 2012; Deorowicz and Grabowski, 2011; Hach et al., 2012; Jones et al., 2012; Kozanitis et al., 2011; Popitsch and von Haeseler, 2013; Rajarajeswari and Apparao, 2011; Tembe et al., 2010) attempt to compress without appeal to a reference. SCALCE (Hach et al., 2012) is one of the most effective, and works by reordering reads within the FASTA file to boost the compression of general purpose compressors. Our approach is able to achieve on average file sizes that are 35% smaller than SCALCE.

Reference-based schemes require a shared reference to be transmitted between all parties who want to decode the data. Most reference-based schemes (e.g. Campagne et al., 2013; Fritz et al., 2011; Jones et al., 2012; Li et al., 2014) focus on compressing alignments between the reads and a set of reference sequences. As such, they work by compressing BAM files, which are the result of alignment tools such as Bowtie (Langmead et al., 2009). The most-used such tool is CRAM (Fritz et al., 2011), which works by mapping reads to a reference sequence and storing any differences. The limitation of these approaches is that they must encode information in the BAM file that can be recreated by re-running the alignment tool. In fact, such BAM compressors may in fact increase the raw size of the data since all the alignment information must be preserved. Another reference-based compressor, fastqz (Bonfield and Mahoney, 2013), attempts to compress sequences directly using its own alignment scheme without first creating a BAM file.

Here, we present a scheme that lies somewhat in the middle of these two extremes: we exploit a shared reference — a compressed transcriptome — but we do no aligning. The reference serves only to generate a statistical, generative model of reads that is then employed in a fixed-order context, adaptive arithmetic coder. The coder is adaptive in the sense that as reads are encoded, the model is updated in such a way that the decoder can reconstruct the updates without any additional information beyond the initial compressed transcriptome. In this way, if the read set differs significantly from the reference transcriptome, the statistical model will eventually converge on this new distribution, resulting in improved compression. We present a scheme that updates the model in such a way that it is robust to sequencing errors, which are a common source of poor compression. By sitting between pure reference-based compression and *de novo* compression, the path encoding scheme gains flexibility and generality: the same scheme works reasonably well even when the provided reference is a poor match for the sample, but is significantly improved with better shared data.

The arithmetic coder uses a fixed-length context to select a conditional distribution for the following base. This scheme is efficient but has the drawback that at the start of each read, there is insufficient context to apply the model. We solve this problem with a new approach. We encode the starts of all the reads in a single, compact data structure called a bit tree. The bit tree is a general scheme for storing sets of small, fixed length (say, < 30 nucleotides) sequences. It is a simplification of other serial encoding schemes such as S-trees (de Jonge et al., 1994) and sequence multiset encoding (Steinruecken, 2014). The bit tree stores sequences but not their order or the number of times they occur in the set. We reorder reads into a standard order to match that stored by the bit tree, and augment the bit tree with auxiliary information indicating the number of repetitions of each sequence.

Taken together, the bit tree for encoding the read starts and the adaptive, context-aware arithmetic coding for the remainder of the read produce files that are on average less than 66% of the size of those produced by a current state-of-the-art *de novo* encoder, SCALCE (Hach et al., 2012). Our approach also produces smaller files compared with reference-based schemes. Its files are on average 33% the size of those produced by CRAM (Fritz et al., 2011) and on average 59% the size of those produced by fastqz (Bonfield and Mahoney, 2013). These are very large improvements in compression, a field where improvements of several percent are often difficult to achieve.

We call the resulting approach *path encoding* because, in the methods section below, we draw a parallel between the design of our arithmetic coder and the problem of efficiently encoding paths in directed graphs, which is a problem that arises in genome assembly (Pevzner et al., 2001) and metagenomic analyses (Iqbal et al., 2012). The bit tree scheme for storing sets of short sequences (kmers) is of independent interest as the need to transmit and store collections of kmers is also increasingly common in de Bruijn-graph-based genome assembly, metagenomic classification (Wood and Salzberg, 2014), and other analyses (Patro et al., 2014).

## 2. RESULTS

### 2.1 Path encoding effectively compresses RNA-seq reads

We selected 7 short-read, RNA-seq data sets of various read lengths and number of reads (Table 1). Both single- and paired-end protocols are represented among the sets. One data set, SRR037452, was chosen because it is a benchmark data set for comparing RNA-seq abundance estimation approaches (e.g. Patro et al., 2014; Roberts and Pachter, 2013). Three sets related to human embryo development were chosen as a representative set of related experiments that one might consider when investigating a particular biological question. A fifth set represents a larger collection of single-end reads of a human cell line. Finally, to assess the effect of using a human transcriptome as a reference when encoding other species, RNA-seq experiments from *Mus musculus* and the bacterium *Pseudomonas aeruginosa* were included. Taken together, these are representative set of RNA-seq read sets.

**Table l:**
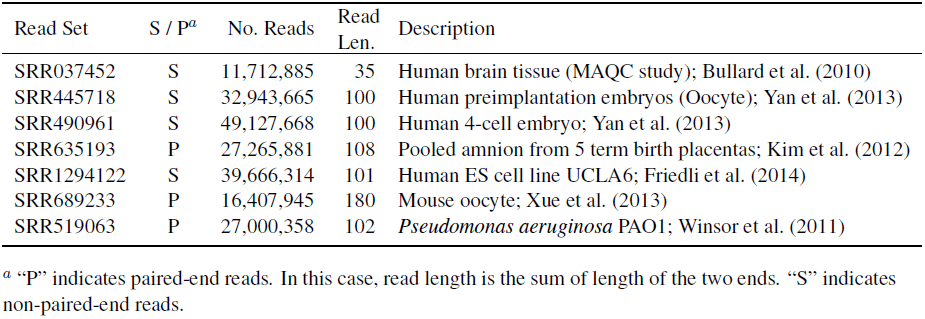
Short-read, RNA-seq data sets compressed in this study.

| Read Set | S / P<sup>a</sup> | No. Reads | Read Len. | Description |
| --- | --- | --- | --- | --- |
| SRR037452 | S | 11,712,885 | 35 | Human brain tissue (MAQC study); Bullard et al. (2010) |
| SRR445718 | S | 32,943,665 | 100 | Human preimplantation embryos (Oocyte); Yan et al. (2013) |
| SRR490961 | S | 49,127,668 | 100 | Human 4-cell embryo; Yan et al. (2013) |
| SRR635193 | P | 27,265,881 | 108 | Pooled amnion from 5 term birth placentas; Kim et al. (2012) |
| SRR1294122 | S | 39,666,314 | 101 | Human ES cell line UCLA6; Friedli et al. (2014) |
| SRR689233 | P | 16,407,945 | 180 | Mouse oocyte; Xue et al. (2013) |
| SRR519063 | P | 27,000,358 | 102 | *Pseudomonas aeruginosa* PAO1; Winsor et al. (2011) |

<sup>a</sup> “P” indicates paired-end reads. In this case, read length is the sum of length of the two ends. “S” indicates non-paired-end reads.

Path encoding is able to reduce these files to 12% – 42% of the size that would be achieved if the file was naively encoded by representing each base with 2 bits (Table 2). The two-bit encoding is approximately 1/4th the size of the sequence data represented as ASCII. (It is approximate because the ASCII encoding includes newline characters separating the reads.) Thus, path encoding reduces files to 3% – 10.5% of the original, raw ASCII encoding. For the human data sets, this is on average 34% smaller than the encoding produced by the SCALCE compression scheme (Hach et al., 2012), a recent, highly effective *de novo* compression approach. This is also smaller than the *de novo* compressor fqzcomp (Bonfield and Mahoney, 2013), which produces files that are larger than those produced by SCALCE.

**Table 2:** Compressed sizes using various methods.

| Read Set | 2-Bit<sup>b</sup> | SCALCE<sup>c</sup> | fastqz<sup>d</sup> | CRAM<sup>e</sup> | PathEnc | No. Trans.<sup>f</sup> |
| --- | --- | --- | --- | --- | --- | --- |
| SRR037452 | 102,487,744 | 66,630,706 | 80,465,928 | 156,554,323 | **43,009,811** | 4.08 |
| SRR445718 | 823,591,625 | 252,989,168 | 238,180,853 | 375,901,891 | **154,873,354** | 0.98 |
| SRR490961 | 1,228,191,700 | 300,176,711 | 316,478,709 | 518,183,711 | **170,546,140** | 0.74 |
| SRR635193 | 736,178,787 | 294,524,283 | 272,862,515 | 366,789,369 | **187,169,742** | 0.90 |
| SRR1294122 | 1,001,574,429 | 299,329,267 | 285,710,714 | 369,774,561 | **187,637,194** | 0.86 |
| SRR689233 | 738,357,525 | 233,812,737 | 266,126,542 | 929,644,204 | **167,586,546** | 1.48 |
| SRR519063 | 688,509,129 | 100,403,786 | 183,275,880 | 714,193,963 | **84,683,932** | 6.14 |

<sup>b</sup> “2-Bit” gives the size of the sequence file in bytes if it were simply encoded using 2-bits per base.
<sup>c</sup> The size of the sequence portion of the SCALCE encoding.
<sup>d</sup> fastqz is a reference-based approach that was provided the same transcriptome as PathEnc.
<sup>e</sup> See Methods for how CRAM’s BAM encoding was adapted to sequence encoding.
<sup>f</sup> The number of transmissions of similar files to make transmitting the shared reference more effective than encoding with SCALCE.

When encoding these files, a human reference transcriptome derived from the hg19 build of human genome was used to prime the statistical model. This transcriptome contains 214,294 transcripts, and occupies 96,446,089 bytes as a gzipped FASTA file. This reference file is required to decompress any files that were compressed using it, but because the same reference transcriptome can be used by many RNA-seq experiments, the cost of transmitting the reference can be amortized over the many files encoded with it. The cost of transmitting or storing the 92 Mb of the reference can be recovered after < 1 to 6.2 transmissions of a compressed file (Table 2, last column). Often the size of the reference transcriptome plus the encoded file is less than the size required by previous pure *de novo* compression schemes — this means that path encoding can also be seen as a very effective *de novo* encoder if the reference is always transmitted with the file, with the option to become a reference-based compressor if several files share the same reference.

Even though this reference contains only human transcripts, it is still effective when encoding RNA-seq experiments from other organisms. For mouse data (SRR689233), for example, the compression is on par with that achieved for the human data sets. For the very different bacterium *Pseudomonas aeruginosa* (SRR519063), the compression gain over *de novo* encoding is still substantial (Table 2, last row). Thus, a single reference can provide enough information to effectively encode RNA-seq reads from many organisms, further allowing its size to be amortized across collections of read sets.

When compared against a recent reference-based encoding scheme, fastqz (Bonfield and Mahoney, 2013), path encoding fares well, consistently producing smaller files than the mapping approach taken by fastqz. In contrast to path encoding, using a mismatched reference for fastqz results in files that are larger than if no reference were used at all (Table 2, last 2 rows). This is because nearly no reads map sufficiently well to the reference. This shows that the non-mapping-based reference scheme implemented by path encoding is both more effective and more robust than mapping-based schemes, which require good matches along a read to benefit from the reference and which also spend a lot of their encoding recording the edits between the reference and the mapped read.

Much of the previous work on reference-based compression has focused on compressing alignment BAM files. The archetypical example of this is CRAM (Fritz et al., 2011). BAM files contain more than sequences. They normally include quality values, sequence descriptions, etc. and may contain multiple alignments for each sequence. To fairly compare sequence compression schemes, we generated BAM files with a single, best alignment for each read to the reference transcriptome, and then stripped extraneous fields (including quality values and sequence names) from the resulting BAM file by setting them to the appropriate “empty” value. These streamlined BAM files were then compressed with CRAM (Table 2). In all cases, path encoding produced much smaller file than CRAM. This is not entirely fair to CRAM, since it attempts to preserve all alignment information in the BAM files, and it also allows for random access to records in compressed file, which path encoding does not. However, for raw compression and transmission, path encoding sequences directly is much more effective than compressing a BAM file. Again, when the reference is mismatched to the sequence (Table 2, last two rows), compression of the CRAM mapping-based approach is reduced substantially.

Figure 1 summarizes the compression achieved by the various methods compared here. It presents encoding sizes as fraction of the naive two-bit encoding of the file. Ratios over 1.0 indicate that an increase in file size over the simplistic encoding.

**Figure 1:**
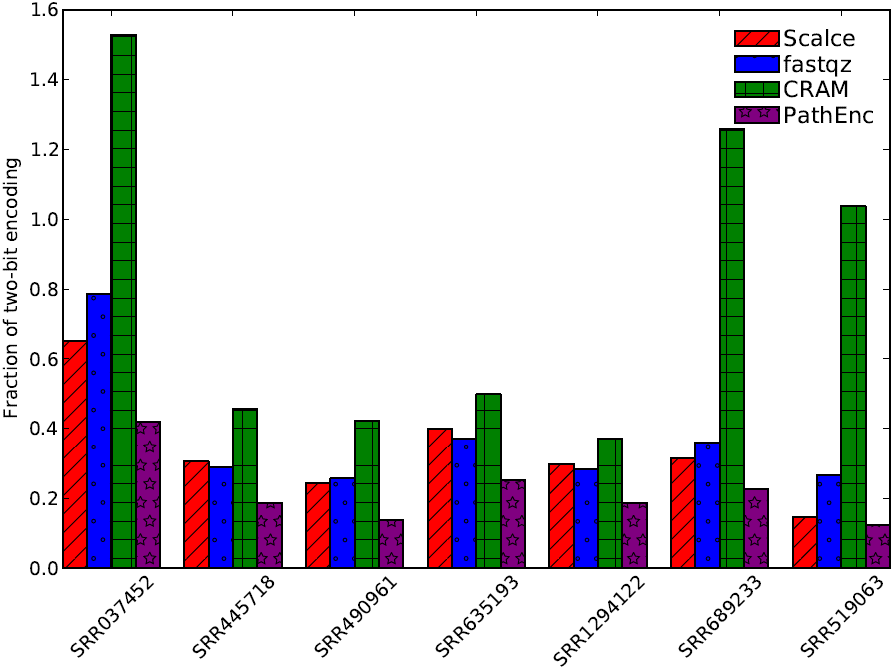
Fraction of two-bit encoding size for files produces by various *de novo* and reference-based methods.

### 2.2 Encoding of the read tails comprises the bulk of the compressed file

A path encoded file consists of several parts (Figure 2). The bulk of the space is used to encode the ends of reads using a context-dependent arithmetic coding scheme (see Methods). This is the part of the encoding that exploits the adaptive statistical model that is initialized from the reference and updated continuously as reads are processed. The first few characters (here 16) of each read are encoded via a bit tree — a data structure that encodes a set of kmers — along with counts for how many reads begin with each kmer (“read head counts” in Figure 2). Together, the read end encoding, the bit tree, and the counts represent the information needed to reconstruct the original reads if we do not care about recording the locations of “N” characters or the orientation of the reads. Since SCALCE and two-bit encoding also do not record the location of “N”s and read orientation is often arbitrary, the sum of the sizes of these three parts are what is reported in Table 2. N locations and the original read orientations can optionally be recovered using the “N locations” and “Flipped bits” parts of the compressed output.

**Figure 2:**
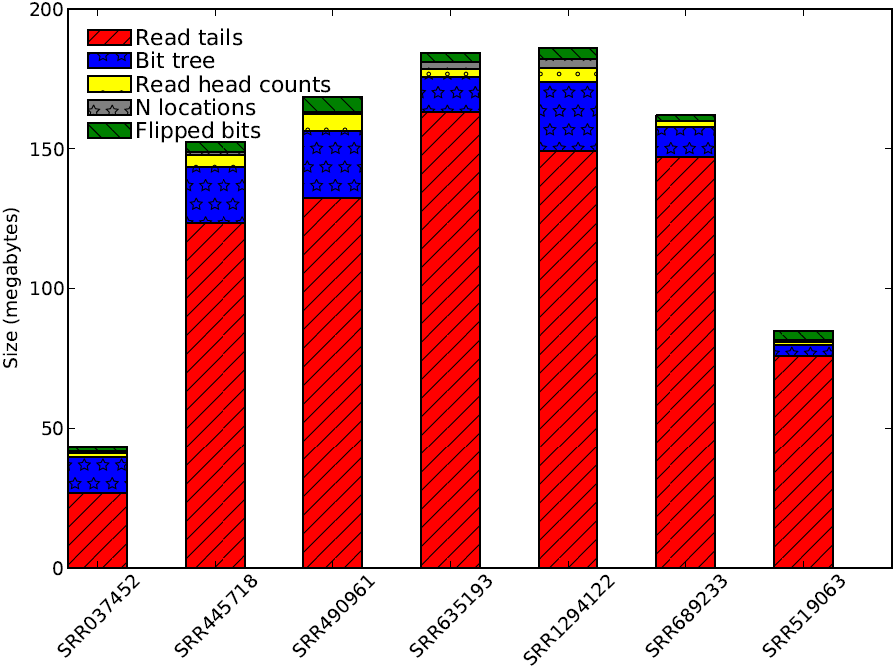
Sizes of the various components of the compressed files. “Read tails” are the portion of the reads encoded using arithmetic encoding. “Bit tree” gives the storage used by the bit tree for encoding the read starts (the first *k* = 16 letters of each read). “Read head counts” is the space taken to store the number of reads with each start. “N locations” is the space to store the location of input Ns that were changed to As upon encoding. “Flipped bits” gives the space needed to record (in a compressed format) a single bit for each read indicating whether the read was reverse complemented.

The read head counts and N locations are compressed simply as gzipped lists of ASCII-encoded, space-separated base-10 integers. The bits recording whether a read was reverse complemented are also simply a gzipped bit vector with a “1” indicating that the read was reverse complemented before it was encoded. While smarter encoding schemes may reduce the size of these parts of the path encoded file (for example by performing a bit-level Burrows-Wheeler transformation (Burrows and Wheeler, 1994) of the bit vector), they do not represent a large fraction of the output and so improvements to them will likely have a small effect.

### 2.3 Priming the statistical model results in improved compression

The availability of the reference typically results in a 15% – 25% reduction in file size for human read sets, a non-trivial gain in compression (Figure 3A). For example, for a file with 3.8 gigabases of sequence (SRR1294122), path encoding with the reference produces an encoded file of 0.17 gigabytes, while starting with a uniform, empty statistical model produces a file of 0.22 gigabytes. For non-human data, the gain of using a human reference is naturally smaller. For mouse reads, the reference yields only a ≈ 3% gain in compression — still a non-trivial size reduction in the context of large files, but much smaller. For the bacterial data, the reference provides little help, but does not hurt compression.

**Figure 3:**
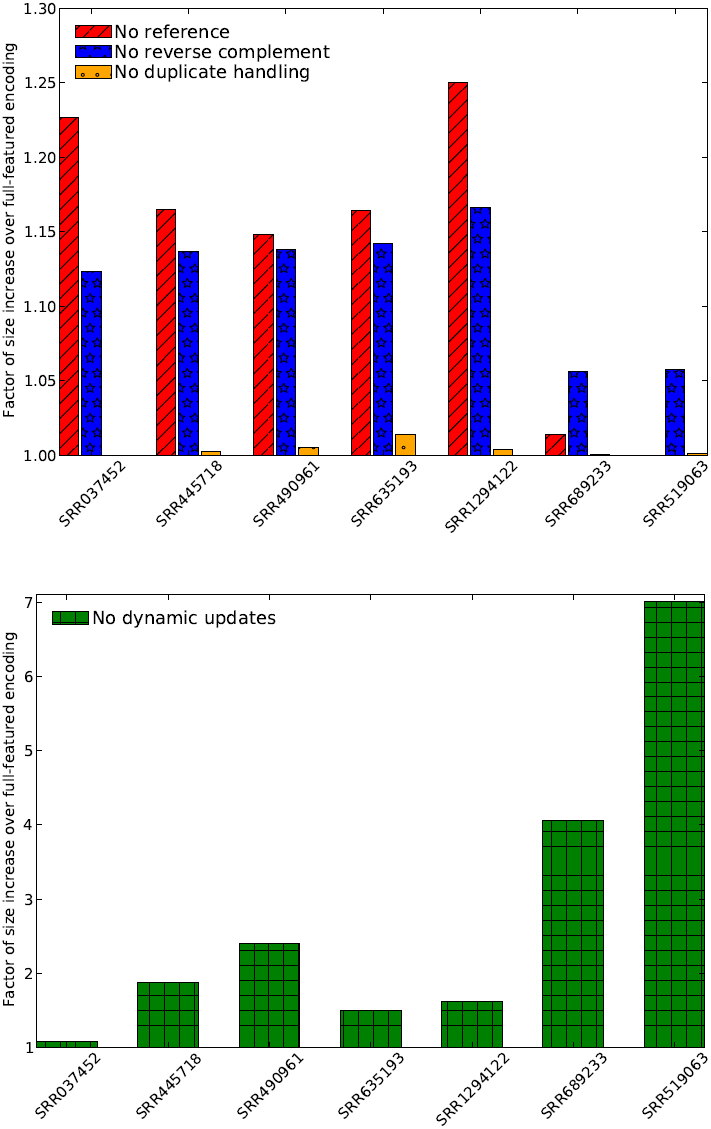
Performance when several features of the path encoding scheme are disabled. All values are given as percentage over the encoding size for the encoding that uses all the features. (A) “No reference” starts with an empty transcriptome reference. “No reverse complement” disables the reverse complementation of the reads. “No duplicate handling” disables the recognition and special encoding of exact duplicate reads. (B) “No dynamic updates” gives the compression when the probabilities of the statistical model are not updated as reads are encoded.

Although the reference provides a starting point for the statistical model, the arithmetic coding we use is adaptive in the sense that read patterns observed frequently during encoding will become more efficiently encoded as they are observed. By disabling these dynamic updates, we can quantify their benefit (Figure 3B), which is substantial. The dynamic updating for the larger human files results in a encodings that are 45% – 92% the size of those produced by the non-dynamic model. For non-human data, where the initial model is likely to be most wrong, the adaptive coding is essential for good compression, resulting in a file that is 4.1 (mouse) or 7.0 (*P aeruginosa*) times smaller. Thus, even with a poor initial model, a good model can be constructed on the fly by adapting the probabilities to the reads as they are processed.

These results show that path encoding provides a unified framework for good compression: when a good reference is available, it can be exploited to gain substantially in compression. When a reference is mismatched to the reads being encoded, the initial model is poor but can be improved via adaptive updates.

### 2.4 The effect of heuristics for reverse complementation and encoding duplicate reads, and of the choice of context size

Reverse complementing reads also provides a significant gain, particularly for reads that match the reference (Figure 3A). This is because the reverse complementation allows the read to agree more with the statistical model. Even when the initial model is poor, reverse complementation is still useful since reads will be reverse complemented to better agree with the model. Recognizing some duplicate reads also leads to a modest improvement in encoding size (Figure 3A). The improvement based on handling duplicate reads is small both because there are relatively few exact duplicate reads and because — in the interest of speed — we only tag a read as a duplicate if every read with the same first 16 bases is identical. It is possible that, in more redundant read sets, better handling of duplicate reads could result in a bigger gains.

Path encoding has one major parameter: the kmer length *k* used to construct the nodes of the context graph (see Methods). A bigger *k* uses more of the preceding string as context to set the probability distribution for the next base, but at the same time bigger *k*s make the effect of sequencing errors last longer since the sequence error affects the context for *k* bases. In addition, a larger *k* requires more memory resources to encode and decode. We find that *k* = 16 is the point at which encoding is most effective (Figure 4). This is also the point at which a kmer can fit in a single 32-bit computer word, leading to an efficient use of memory. While longer *k* does reduce the size of the encoding of the read ends (both because the read ends are shorter and because a longer context is used), the size of the bit tree encoding the read starts grows more quickly than the savings gained.

**Figure 4:**
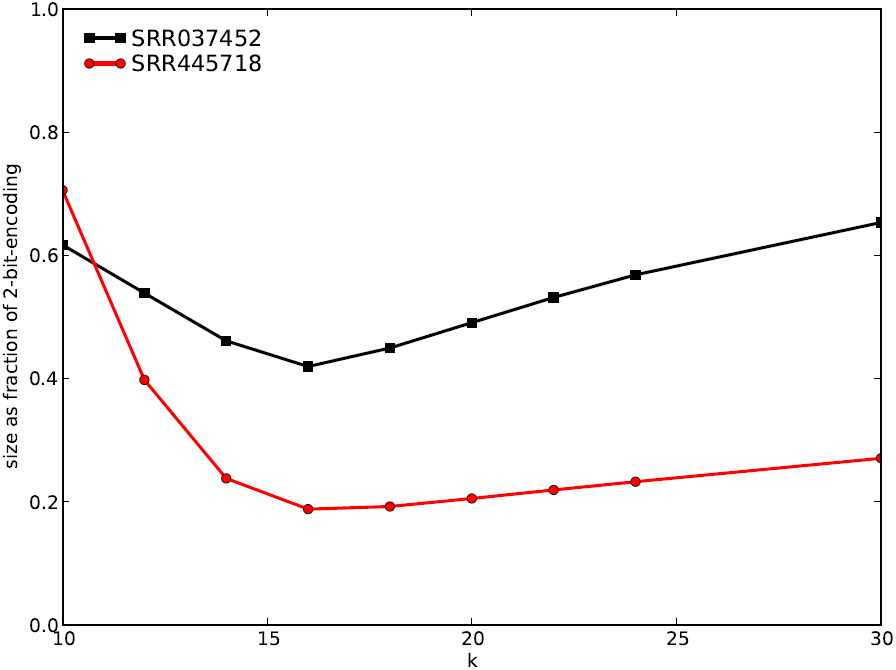
File size, represented as a fraction of the two-bit encoding size, using various kmer lengths *k*.

### 2.5 Encoding and decoding path encoded files is fast

Path encoding the entire data set here takes 2.37 hours. This is nearly identical to the running time for running bowtie and CRAM on the same data set. Times for encoding individual files ranged from 5.1 minutes to 32 minutes. Decoding is generally much faster, never taking longer than 22 minutes for the files in Table 1. Since encoding is typically performed only once, and decoding is performed multiple times, this is the trade-off that is normally desired: very effective compression, taking longer to encode, with quick decoding.

**Figure 5:**
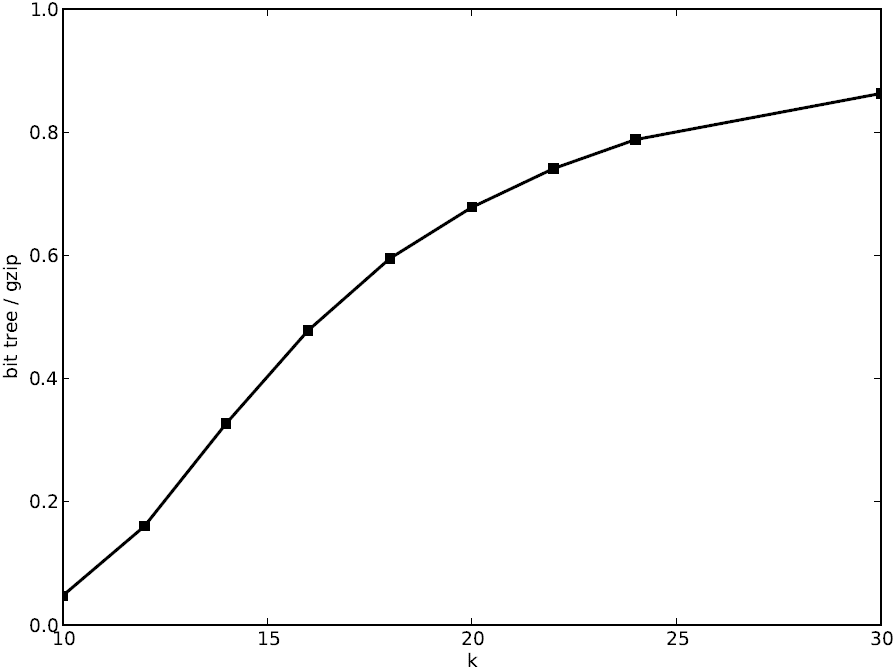
Size of the bit tree for storing sets of kmers of various lengths (*k*). Each kmer set is the first *k* characters of each read in SRR037452, and the size is plotted as fraction of the size obtained for compressing the same set using gzip -9.

## 3. DISCUSSION

We have provided a novel encoding scheme for short read sequence data that is effective at compressing sequences to 15% – 25% of the uncompressed, two-bit encoded size. To do this, we introduced the novel approach of encoding paths in a de Bruijn graph using an adaptive arithmetic encoder combined with a bit tree data structure to encode start nodes (see Methods for a description). These two computational approaches are of interest in other settings as well. Path encoding achieves better compression than both *de novo* schemes and mapping-based reference schemes. Since the reference for the human transcriptome is small (92 Mb) compared with the size of the compressed files, the overhead of transmitting the reference is recovered after only a few transmissions. In addition, the reference is merely a gzipped version of the transcriptome — a file that most researchers would have stored anyway.

Path encoding is also more general than reference-based schemes since we have more flexibility in choosing how to initialize the statistical model with the reference sequence. For example, the reference could be reduced to simple context-specific estimates of GC content. This will naturally lead to worse compression but will also eliminate most of the need to transmit a reference. Technology-specific error models could also be incorporated to augment the reference to better deal with sequencing errors. In addition, SNP data from a resource such as the HapMap project could be included in the reference to better deal with genomic variation. Framing the problem using a statistical generative model as we have done here opens the door to more sophisticated models being developed and incorporated.

Another source of flexibility is the possibility of lossy sequence compression. While encoding, a base that has low probability in a particular context could be converted to a higher probability base under the assumption that the low probability base is a sequencing error. Implementation of this technique does indeed reduce file sizes substantially, but of course at the loss of being able to reconstruct the input sequence. While lossy compression may be appropriate for some analyses (such as isoform expression estimation) and error correction can be viewed as a type of lossy encoding, because we are interested in lossless compression, we do not explore this idea more here.

A recent line of work (e.g. Daniels et al., 2013; Janin et al., 2014; Loh et al., 2012) aims at producing searchable, compressed representations of sequence information. Allowing sequence search limits the type and amount of compression that can be applied and requires some type of random access into the encoded sequences. Arithmetic encoding does not generally allow such random access decoding because the constructed interval for a given symbol depends on all previously observed symbols. However, decompression with our path encoding scheme can be performed in a streaming manner: the encoded file is read once from start to finish, and the decoder produces reads as they are decoded. This would allow reads to be decoded as they are being downloaded from a central repository.

The other dimension of compressing short-read data is storing the quality values that typically accompany the reads. Path encoding does not attempt to store these quality values as there are other, more appropriate approaches for this problem (Cánovas et al., 2014; Hach et al., 2012; Ochoa et al., 2013; Yu et al., 2014). Path encoding can be coupled with one of these approaches to store both sequence and quality values. In fact, in many cases, the quality values are unnecessary and many genomic tools such as BWA (Li and Durbin, 2009) and Sailfish (Patro et al., 2014) now routinely ignore them. Yu et al. (2014) showed that quality values can be aggressively discarded and without loss of ability to distinguish sequencing errors from novel SNPs. Thus, the problem of compression of quality values is both very different and less important than that of recording the sequence reads.

An open-source implementation of path encoding is available from http://www.cs.cmu.edu/∼ckingsf/ software/pathenc/ and as supplementary material.

## 4. METHODS

### 4.1 Overview

Our compression approach is composed of several different encoding techniques that are applied to the input reads as a set. First, the reads are reverse complemented based on a heuristic to determine which orientation matches the initial reference better (Section 4.6). The initial *k* letters of each read are stored in a bit tree data structure along with the counts of their occurrences (Section 4.3). These initial *k* letters of each read are called the read *head*. The reads are then reordered to place reads with the same heads next to one another. Finally, the remainder of each read (called the read *tail*) is encoded using an adaptive arithmetic coding scheme (Section 4.4) inspired by the path encoding problem (Section 4.2).

### 4.2 The path encoding problem

We can capture much of the information in a reference transcriptome using a graph *G* that has a node for every kmer that occurs in a transcript and an edge (*u, v*) between any two kmers *u* and *v* if *v* follows *u*, overlapping it by *k* — 1 characters, in some transcript. This is a de Bruijn graph, except it is derived from several strings rather than a single string. A read *r*, if its sequence occurs in the transcriptome, corresponds to a path in *G*, and conversely there is only one path in *G* that spells out *r*. Therefore, *r* can be encoded by specifying a path in *G* by listing a sequence of nodes. This leads to a very general problem:

> **Problem** (Path encoding). *Given a directed graph G, encode a collection of paths P*_1_*,P*_2_*,…, P_n_, each given as an ordered sequence of nodes of G, using as little space as possible*.

Our compression scheme uses one system for encoding the first node of each read path *P_i_* (the read head) and another system for encoding the remaining nodes in the path (the read tails). The first system is based on encoding a depth-first search of a tree that represents the first nodes of each path. The second system is based on adaptive, context-aware arithmetic coding. The challenge posed by adaptive arithmetic coding in this context is to not be mislead into assigning large probability to sequencing errors. We describe each approach below.

### 4.3 Encoding the starts of the reads with a bit tree

Let *T* be the kmer trie defined as follows. *T* has a root node that has four children, and each edge from the root node to a child is labeled by a different nucleotide in *{A, C, G, T*}. Each of these children themselves have four similar children, with edges for each of the nucleotides. This continues until every path from the root to a leaf node has exactly *k* edges on it. In this way, a complete, 4-ary tree of depth *k* is constructed such that any path from the root to a leaf spells out a unique kmer, and every possible kmer corresponds to some such path in *T*. The set of kmers *K* that appear at the start of some read corresponds to a subset of those possible paths, and we can construct a subtree *T*_|*K*_ from *T* by removing all edges that are not used when spelling out any kmer in *K*. Knowing *T*_|*K*_ allows us to reconstruct *K* precisely: *K* consists of those kmers spelled out by some path from the root to a leaf in *T*_|*K*_.

*T*_|*K*_ can be encoded compactly by performing a depth-first search starting at the root, visiting each child of every node in a fixed order (say A then C then T then G) and emitting a 1 bit whenever an edge is traversed for the first time and a 0 bit if we attempt to go to a child that does not exist. This bit stream is then compressed using a general purpose compressor (gzip; Gailly and Adler, 2014). *T*_|*K*_ can be reconstructed from this stream of 0s and 1s by performing a depth-first search on *T* traversing a edge whenever a 1 bit encountered but pruning subtrees whenever a 0 bit is read. *K* is then reconstructed as the set of kmers corresponding to the leaves we encountered.

The trie *T* never need be actually built to perform the encoding or the decoding. Rather, a sorted list of the kmers is sufficient for simulating the traversal of the trie to encode, and decoding only ever needs to implicitly construct the part of the trie that is on the current depth-first search path. Pseudocode for bit tree encoding is given in Algorithm 1. In practice, encoding and decoding of very large collections of kmers takes very little time or memory.

**Algorithm 1.**
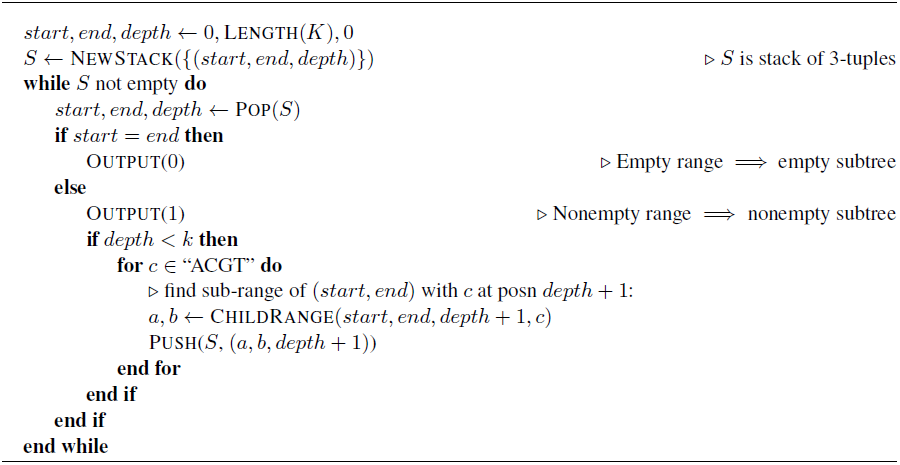
Algorithm to encode set of kmers *K* using a bit tree.

The same kmer may start many reads, but the encoding of *T*_|*K*_ only records which kmers were used, not the number of times each was used. To store this, we write out a separate file called the count file with the count of each kmer in *T*_|*K*_ in the order that the kmers will be visited during the decoding of *T*_|*K*_. This file stores counts as space-separated ASCII integers, and the entire file is compressed using the gzip (Gailly and Adler, 2014) algorithm. *T*_|*K*_ also does not record the order in which the kmers were used as read heads, so we reorder the read set to put reads with the same start adjacent to one another in the same order as their starts will be encountered during the decoding of *T*_|*K*_.

This data structure is essentially the same as a S-tree (de Jonge et al., 1994) specialized to kmer tries, except that no data is stored at the leaves, and because the length of every sequence is a known constant *k* we need not store any information about the (always nonexistent) children of nodes at depth *k*. It is a simplification of Steinruecken (2014) since counts are stored only for the leaves.

### 4.4 Arithmetic coding of read tails

Arithmetic coding (Moffat et al., 1998; Rissanen and Langdon, 1979; Witten et al., 1987) compresses a message by encoding it as a single, high-precision number between 0 and 1. During the encoding, an interval [*a, b*] is maintained. At the start, this interval is [0,1], and at each step of the encoding, it is reduced to a subinterval of the current interval. At the end, a real number within the final interval is chosen to represent entire message. The interval is updated based on the probability of observing each symbol in a particular context. For path encoding, we store a probability distribution *p_u_*(·) associated with each node *u* in *G* on its outgoing edges such that *p_u_*(*v*) gives an estimate for the probability that edge (*u, v*) will be the one used by a path leaving *u*. We also give the outgoing edges of *u* an arbitrary, fixed order 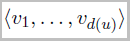, where *d*(*u*) is the out-degree of *u* (when encoding DNA or RNA, *d*(*u*) is always 4). Using this ordering, we can compute the cumulative distribution 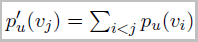. The probability distributions *p_u_*(·) for each node in *G* represent the statistical generative model that encodes the information about which sequences are more or less likely.

Let 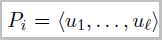 be a sequence of nodes of the path *P_i_* that we are encoding. When beginning to encode *P_i_*, the current node is *u*_1_, the current interval is [0,1]. The first node *u*_1_ is encoded using the bit tree approach described above. Suppose we have encoded *u*_1_,…, *u*_*j*-1_ and the current interval is [*a, b*]. To encode *u_j_*, we update the interval to:

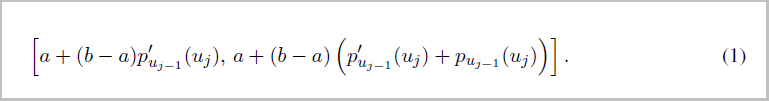

This chooses a subinterval of [*a, b*] that corresponds to the interval for *u_j_* in 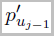. The intuition for why this approach achieves compression is that it requires fewer bits to specify a number that falls in a high-probability (large) interval than in a low probability (small) interval. Therefore, if we choose the distributions *p_u_*(·) well so that common edges are given high probability, we will use few bits to encode frequently occurring symbols.

In practice, Equation 1 is not used directly because it would require infinite precision, real arithmetic which is not available on digital computers. Rather, an approach (Moffat et al., 1998) that uses only finite, small precision, integer arithmetic and that rescales the current interval when necessary is used. This practical arithmetic coding has achieved state-of-the-art compression in many applications (Rissanen and Langdon, 1979; Witten et al., 1987).

### 4.5 Initializing and updating the sequence generative model

The probability distributions *p_u_*(·) for each node *u* specify what we consider to be a high-probability sequence (which is equivalent to a high probability path). It is here that we can use shared, prior information to influence the encoding. These distributions need not be constant — so long as the decoder can reconstruct any changes made to the distributions, we can adapt them to observed data as we see it.

We derive *p_u_(v)* using counts *c_u_*(*v*), which are set according to:

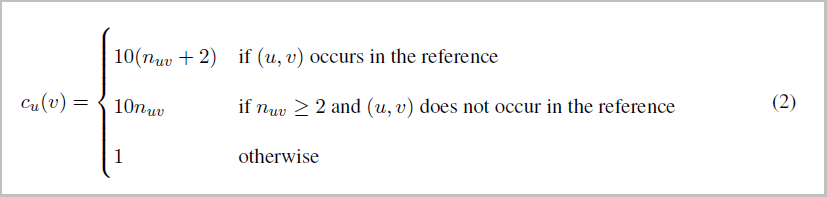

where *n_uv_* is the number of times edge (*u, v*) was observed in the read paths that have been encoded so far. This expression for *c_u_*(*v*) requires an edge (*u, v*) to either occur in the reference or be used at least twice in a read path before it is given the larger weight. This is to reduce the impact of sequencing errors that are frequent, but unlikely to occur twice. The 1 in the third case of Equation 2 acts as a pseudocount for edges that have not yet been observed, and the 10 in the first two cases sets the relative weight of observations versus this pseudocount. We compute *p_u_*(*v*) *= c_u_*(*v*)/Σ_*w*_*c_u_*(*w*).

During the encoding of reads, it is possible that we encounter a kmer that we have never seen before. In this case, we encode the base following this kmer using a default probability distribution derived from a distinct count distribution *c_0_*(*b*) that gives the number of times we encoded base *b* using this default distribution. After the first time we see a new kmer, we add it to the graph *G* and on subsequent observations, we treat it using Equation 2.

An alternative way to view the arithmetic coding scheme above is that the probability distribution is provided by a fixed-order (*k*) Markov chain for which the transition probabilities are updated as edges are traversed. The *k* preceding bases provide a context for estimating the probability of the next base. The read heads provide the initial context for the Markov chain. This view also motivates the need to handle sequence errors and (less common) sequence variants effectively, since an error in a read will produce an incorrect context for *k* bases, resulting in decreased compression. This is the reason for the step-function update rule in Equation 2. We found that using the read head as the context at the start of the read, rather than concatenating all the reads together, resulted in improved compression because it was no longer necessary to encode bases using a context that consisted of the non-biological joining of two reads.

### 4.6 Other considerations

The reference model only includes the forward strand of the transcript but, in an unstranded RNA-seq protocol, reads may come from either strand. We implement a heuristic for selecting whether or not to reverse complement the input read. To do this, we estimate whether the forward read *r* or its reverse complement *rc*(*r*) will produce a smaller encoding by summing the number of observations of the kmer transitions in the read that also occurred in some reference transcript for both *r* and *rc*(*r*). If *rc*(*r*) has a higher number of observed transitions in the statistical model, we reverse complement the read before encoding. The decision to reverse complement reads is made for all reads at the beginning of compression before any reads are encoded and before the read starts are encoded. We typically do not need to store whether a read was flipped or not since the choice of strand recorded in the file was arbitrary to begin with. However, if it is important to store the string in the direction it was originally specified, a single bit per read is recorded indicating whether the read was reverse complemented.

Due to biases in RNA-seq, and due to pooling of technical replicates, it is often the case that the exact same read sequence is listed more than once in a read file. To more compactly encode this situation, we check whether the set of reads that start with a given kmer *m* consists entirely of *d* duplicates of the same sequence. In this case, we record the number of reads associated with *m* in the count file as —*d* rather than *d*, and we only store one of the tails in the path file. Encoding of duplicates in this way provides a small decrease in size for most files, but when many duplicate reads are present can result in a large decrease.

To simplify the statistical model, any Ns that appear in the input file are translated to As upon initial input. This is a strategy taken by other compressors (Hach et al., 2012) because the lowest quality value always indicates an N and all Ns must have the lowest quality value. If quality scores are not stored with the sequences and the locations of the Ns are needed, a separate file is output with their locations.

Because reads are reordered, the two ends of a mate pair cannot be encoded separately if pairing information is to be preserved. Instead, when dealing with paired-ended RNA-seq, we merge the ends of the mate pair into a longer read, encoding this “read” as described above. If the library was constructed with the mate pairs from opposite strands, one strand is reverse complemented before merging so that the entire sequence comes from the same strand in order to better match the generative model described above. This transformation can be undone when the reads are decoded.

### 4.7 Implementation details

Software implementing the path encoding and decoding method was written in the Go programming language, using a translation of the arithmetic coding functions of Moffat et al. (1998). The software is parallelized and can use up to 8 threads to complete various steps of the encoding and decoding algorithms simultaneously. To limit memory usage, the counts *c_u_*(·) described above were stored in 16-bit fields. This resulted in some loss of compression effectiveness compared with 32-bit fields but large improvements in running times for the larger files. The software is open source and freely available at http://www.cs.cmu.edu/∼ckingsf/software/pathenc.

### 4.8 Comparison with other methods

SCALCE (Hach et al., 2012) version 2.7 was run with its default parameters, using `−r` for paired-end read sets. The file sizes reported are the sizes of the `.scalcer` files it produces, which encode the sequence data (except the positions of the Ns). The program fqzcomp (Bonfield and Mahoney, 2013) version 4.6 was run using the recommended parameters for Illumina data (`−n2 −s7+ −b −q3`). The file sizes used were the sizes of only the portion of its output file that encodes for the sequences, as printed by fqzcomp. Running fqzcomp with `−s8` instead of `−s7+` produced files that were still larger than SCALCE. For paired-end reads, fqzcomp often achieved better compression if it was provided with a FASTQ file that contained both ends merged into a single read, and so sizes for compressing these files (the same as provided to path encode) were used. Despite this, fqzcomp always produced files that were larger than SCALCE, and so only the SCALCE numbers are reported. Both SCALCE and fqzcomp are *de novo* compressors. Experiments with the *de novo* version of fastqz always produced larger files than fqzcomp and so its results are not reported here. The reference-based version of fastqz (Bonfield and Mahoney, 2013) version 1.5 was provided the same reference as used with path encoding (a multi-fasta file with transcripts), processed with the `fapack` program. The file sizes reported for fastqz are the sum of the sizes of its output files `.fxb.zpaq` and `.fxa.zpaq` that encode the sequences (except the Ns).

CRAM (Fritz et al., 2011) is designed for compressing BAM files. To adapt it to compress sequences, read files were aligned with Bowtie (Langmead et al., 2009) using `––best −q −y ––sam` to an index built from the same transcriptome as used for path encoding. Quality values, sequence names, and sequence descriptions were stripped from the file (fields 1 and 11), MAPQ values were all set to 255, and the RNEXT and PNEXT fields were set to “*” and “0” respectively. The resulting simplified SAM file was converted to a sorted BAM file using samtools (Li et al., 2009). This file was then encoded using CRAM, and the reported file size is that of the resulting `.cram` file. (Leaving the MAPQ, RNEXT and PNEXT fields unchanged resulted in compressed files of nearly identical size.)

## 5. ACKNOWLEDGMENTS

This work has been partially funded by the US National Science Foundation (CCF-1256087, CCF-1053918) and US National Institutes of Health (R21HG006913 and R01HG007104). C.K. received support as an Alfred P. Sloan Research Fellow. We would like to thank Darya Filippova, Emre Sefer, and Hao Wang for useful discussions relating to this work and for comments on the manuscript.

